# Emergent collective organization of bone cells in complex curvature fields

**DOI:** 10.1101/2020.10.28.358572

**Authors:** Sebastien J.P. Callens, Daniel Fan, Ingmar A.J. van Hengel, Michelle Minneboo, Lidy E. Fratila-Apachitei, Amir A. Zadpoor

## Abstract

Individual cells and multicellular systems have been shown to respond to cell-scale curvatures in their environments, guiding migration, orientation, and tissue formation. However, it remains unclear how cells collectively explore and pattern complex landscapes with curvature gradients across the Euclidean and non-Euclidean spectra, partly owing to fabrication limitations and the lack of formal geometric considerations. Here, we show that micro-engineered substrates with controlled curvature variations induce the collective spatiotemporal organization of preosteoblasts. By leveraging mathematical surface design and a high-resolution free-form fabrication process, we exposed cells to a broad yet controlled, heterogeneous spectrum of curvature fields. We quantified curvature-induced spatial patterning at different time points and found that cells generally prefer regions with at least one negative principal curvature. We also show that multicellular cooperation enables cells to venture into unfavourably-curved territories, bridging large portions of the substrates, and collectively aligning their stress fibres. We demonstrate that this behaviour is partly regulated by cellular contractility and extracellular matrix development, underscoring the mechanical nature of curvature guidance. Our findings offer unifying perspectives on cell-geometry interactions that could be harnessed in the design of micro-engineered biomaterials, for example, for tissue engineering applications.

## 1. Introduction

The dynamic, bidirectional interactions between cells and their intricate environment orchestrate tissue morphogenesis, homeostasis, and repair, and are implicated in numerous diseases ^1–3^. The complexity of the extracellular environment is not only due to its diverse and heterogeneous composition but is also caused by its hierarchical spatial structure that imposes geometrical constraints on the force-generating cells ^4,5^. Cells have long been known to sense such geometrical cues at subcellular scales ^6,7^, yet recent evidence shows that geometrical features at much larger scales also affect cell migration, differentiation, and fate, as well as tissue shape and growth kinetics ^8^. Unravelling this interplay between cells and the shape of their surroundings is key to advance the design of artificial scaffolds and biomaterials, where geometry can be harnessed as a micro-engineered cell cue ^9–11^.

From a mathematical viewpoint, the local geometry of the extracellular environment can be fundamentally characterized using the concept of surface curvature ^12^. In recent years, numerous studies have begun to address the role of cell-scale curvature on the dynamics and organisation of cells and tissues. Indeed, curvature guidance has been observed in the directional migration and preferential orientation of a variety of individual cells and multicellular monolayers ^13–17^. Moreover, various cell types have expressed an overall preference for local concavities as opposed to convexities ^18,19^. Biophysical models have suggested key roles for cytoskeletal contractility and nuclear deformation in this large-scale curvature sensation, generally implying that cells with pronounced stress fibres avoid bending and search for relaxed configurations ^20–23^. Despite the availability of such pioneering findings, it remains elusive how cells behave in more complex curvature landscapes. Early studies typically resorted to substrates with limited architectural complexity, involving cylindrical wires ^13,24,25^ or patterned hemispherical substrates ^26,27^, precluding many physiologically-relevant geometries, including saddle shapes or sharp curvature transitions. Moreover, the mathematical descriptions of the surface curvature have not received much attention, hampering the development of a unified, unambiguous theory of cell-scale curvature guidance. Indeed, many studies have considered only a single class of curved substrates ^24,25^, or have relied exclusively on the concepts of convexity and concavity instead of the fundamental definitions of curvature as described by differential geometry ^19,26^.

Here, we adopt a geometry-centred perspective and demonstrate collective spatiotemporal cell organization in precise environments with varying curvature distributions. To this end, we designed several substrates, derived from mathematically-defined surface families, covering a wide range of cell-scale types of curvature variation. Using high-resolution multiphoton lithography (*i.e.*, a 3D printing technique with submicron resolution) and replica moulding, we fabricated chips on which we cultured murine preosteoblasts for several days. Our focus on bone-like cells was motivated by the ongoing quest for geometrically-optimized biomaterials that enhance bone tissue regeneration. While previous studies have either focused on individual cell behaviour at short time scales ^14,19^ or on tissue-level performance in larger-scale environments ^28–30^, we studied curvature guidance at the intermediate time points where cells collectively pattern their environment and establish a template for bone-like tissue formation. By mapping 3D confocal microscopy data to the underlying curvature distributions, we explored the rules for emergent collective cell patterning. Specifically, we found that the curved-to-planar transitions, often ignored in earlier studies, are highly attractive to cell collectives and that cell sheets spontaneously detach from certain curved regions, thereby altering the extracellular geometry sensed by new cells. Moreover, we studied curvature-guided stress fibre orientation and investigated the important role of contractility in collective curvature guidance. Our results provide unifying perspectives on curvature-driven collective cell organization, paving the way towards the geometric optimization of micro-engineered environments.

## 2. Results

### 2.1. Development of cell substrates with controlled curvatures

We first set out to design substrates that would expose cells to a broad, yet controlled spectrum of curvatures. A complete description of surface curvature requires two independent curvature measures. The most common choices are either the two principal curvatures (*i.e.*, the maximum and minimum curvatures, *κ*_1_ and *κ*_2_, respectively), or the pair of the mean 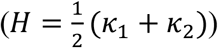 and Gaussian (K = *κ*_1_*κ*_2_) curvatures (Figure 1a). We explored different surface families based on their mean and Gaussian curvature profiles, and focused on axisymmetric surfaces, as these could readily be converted to printable substrates. The first geometry that we selected was the unduloid, which is a simply-periodic surface family with constant, non-zero mean curvature. An unduloid interpolates between a cylinder and a string of connected spheres, depending on its specific parametrization (Supplementary Movie 1) ^31^. This interpolative nature enabled us to select a cylinder, a set of spheres, and an intermediate unduloid all of which with the same (constant) mean, yet different Gaussian curvatures (Figure 1b,e). Next, we selected two saddle surfaces (*i.e.*, *K* < 0): the pseudosphere, having constant negative Gaussian curvature (as opposed to a sphere with constant positive Gaussian curvature), and the catenoid, having constant zero mean curvature (*i.e.*, a minimal surface). Since these surfaces are not simply-periodic by nature, we designed strings of repeating pseudospheres and catenoids, in accordance with the other surfaces (Figure 1b). Finally, we included a sinusoidally-deformed cylinder. In contrast to a normal cylinder (K = *κ*_2_ = 0), this deformed variant is enriched with alternating regions of positive and negative Gaussian curvatures (Figure 1d).

**Figure 1:**
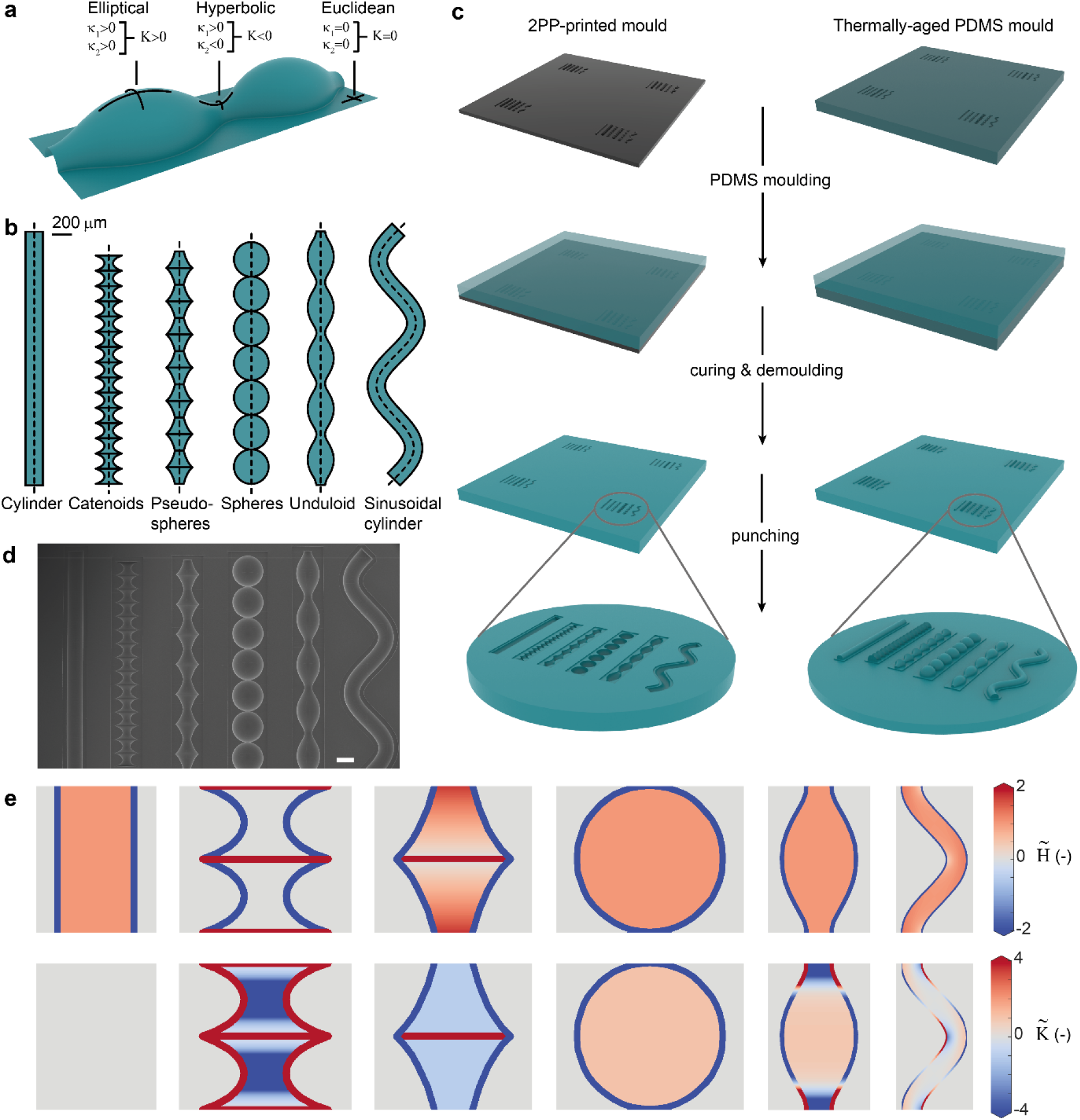
Design and microfabrication of curved cell substrates. a) Local surface geometry defined in terms of the principal curvatures, *κ*_1_ and *κ*_2_, and the Gaussian curvature, *K*. b) The surface profiles (top view) used to design curved cell substrates. The first five surfaces are surfaces of revolution (the dotted line is the rotation axis), while the last surface is obtained by sweeping a circle along a sinusoidal path (dotted line). c) Fabrication of the PDMS substrates with curved imprints (concave, left column) or protrusions (convex, right column). Left: A master-mould containing four sets of curved substrates (protruding halves of the surfaces of revolution) was fabricated using 2PP. Next, PDMS counter-moulding was performed and circular specimens were stamped and prepared for cell culture. Right: A PDMS mould with imprinted substrates was thermally aged and used as a master-mould for moulding specimens with protruded (convex) substrates. The illustration is not shown to scale. d) SEM image of a PDMS sample with imprinted (concave) substrates. Scale bar represents 200 μm. e) Projected curvature maps for the six types of substrates, displaying repetitive unit cells. The top and bottom rows represent the normalized mean and Gaussian curvatures, respectively. From left to right: cylinder, catenoids, pseudospheres, spheres, unduloid, and sinusoidal cylinder. The curvatures are visualized for the convex substrate variants. For the concave substrates, the mean curvatures are equal in magnitude but opposite in sign. The Gaussian curvatures remain the same.

We sized the surfaces to appropriate cell-scale dimensions, based on previous studies ^14,19^, and used them as templates for half-revolution master moulds that were 3D printed using two-photon polymerization (2PP). Single- and two-step replica moulding with poly(dimethylsiloxane) (PDMS) provided us with precisely curved cell culture environments, consisting of both concave imprints (*H* < 0) and convex protrusions (*H* > 0) of the same surfaces (Figure 1c,d). By using both the concave and convex variants, we could significantly expand our total curvature spectrum, as these substrates feature the same Gaussian curvatures, yet opposite mean curvatures.

### 2.2. Murine preosteoblasts prefer regions with negative minimum principal curvature

To investigate curvature-guided spatiotemporal cell patterning, we cultured murine preosteoblasts (MC3T3-E1) on the curved substrates for several days. This cell line has been used before in the context of curvature-driven tissue growth ^30,32^. After 5 days, we observed confluent layers on the planar regions and curvature-dependent patterning on the non-planar regions (Figure 2a and Supplementary Figure 1a). After 8 days, this trend continued and large cell collectives were found to differentially cover the substrates. The frequency maps of the spatial actin distribution, created by superimposing confocal image projections, revealed strong differences in the patterning on the concave (*H* < 0) and convex (*H* > 0) variants of the substrates (Figure 2b and Supplementary Figure 1b-d). Uniform cell coverage was observed in the concave substrates, while the convex variants exhibited distinct regions with high and low actin intensities. On these convex substrates, we found more coverage on the hyperbolic regions (saddle-shaped, *K* < 0) than on the elliptical regions (sphere-like, *K* > 0), as exemplified on the unduloid substrate with a constant mean and varying Gaussian curvatures (Figure 2a,b). On the convex catenoids and pseudospheres (*K* < 0), we observed full cell coverage along the entire substrate, except for the sharp (locally elliptic) transition regions between the saddles. Moreover, we consistently found high cell densities at the transition regions between the convex structures and their planar surroundings, while the reverse was observed at the concave-to-planar transitions. Taken together, these observations demonstrate the collective preference of the cells to pattern regions where the minimum principal curvature is negative (*i.e.*, *κ*_2_ < 0), which includes all regions with at least one concave direction (Figure 2c). This translates to regions with either *K* < 0 (saddle shapes), or K ≥ 0 combined with *H* < 0 (*e.g.*, concave spheres). Indeed, the convex-to-planar transitions, that feature strong cell attraction, are also characterized by *κ*_2_ < 0. This general preference for *κ*_2_ < 0 was confirmed when considering the mean actin intensity across the full curvature spectrum presented to the cells, showing higher intensities for regions with a negative minimum principal curvature (Figure 2d). Nevertheless, the frequency maps and the intensity plots also show that cells do not entirely avoid unfavourably-curved regions. For example, substantial regions of the spherical substrates were covered with cells, despite the constant positive *κ*_2_, suggesting a collective ability to conquer such less favourable curvatures.

**Figure 2:**
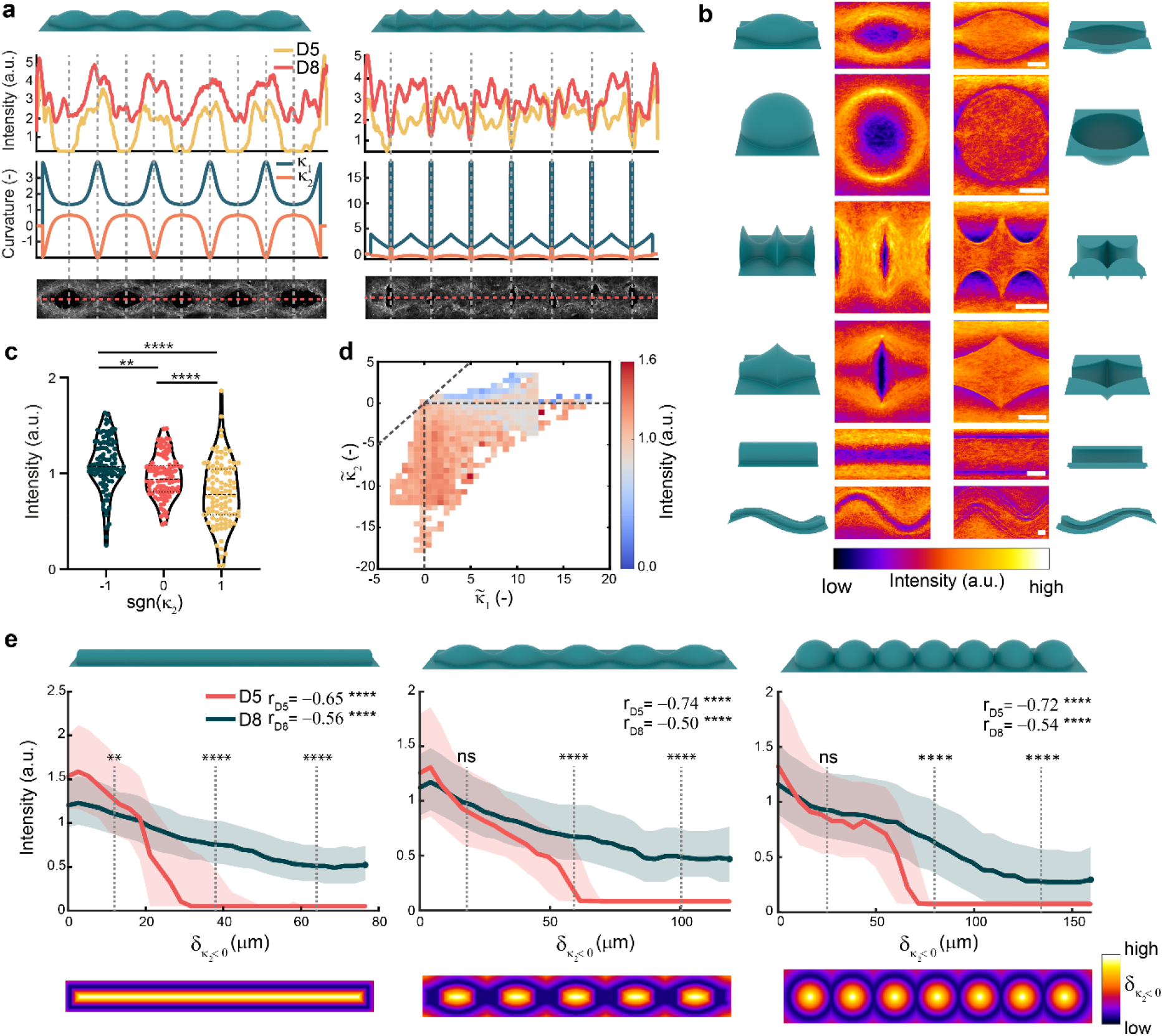
Collective spatiotemporal cell patterning on curved substrates. a) Normalized actin intensity, measured along the centreline, varies as a function of the principal curvatures *κ*_1_ and *κ*_2_. The intensity profiles are obtained from multiple specimens (*n* > 3) for the convex unduloid (left) and convex pseudosphere (right) substrates (see also Supplementary Figure 1a). b) Frequency maps displaying spatial actin patterning on day 8. The data is obtained by stacking periodic units from multiple images (*n* > 15) for both the convex (left column) and concave (right column) variants of the six substrates (see also Supplementary Figure 1b-d). The scale bars represent 100 μm. c) Normalized actin intensity versus the sign of *κ*_2_ for convex substrates on day 8. 100 random data points for each category were sampled from all the available data (superpixels of 80×80 pixels). The data are shown as violin plots with median and interquartile range. The Welch’s ANOVA with Games-Howell’s multiple comparisons test: ** *p* < 0.01, **** *p* < 0.0001. d) Heat map of the median normalized actin intensity *vs.* the two normalized principal substrate curvatures 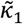 and 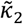 for all substrates on day 8. The data points are obtained by rasterizing the projected confocal images in elements of 20 by 20 pixels. e) Normalized actin intensity reduces with increasing distance to the closest region with 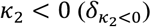. The data are shown for three convex substrates (*n* > 3) and two time points (day 5 and day 8). The solid line represents the median value, while the shaded areas correspond to the interquartile range. The *r*-values represent the Spearman’s correlation coefficients. Mann-Whitney U tests: ** *p* < 0.01, **** *p* < 0.0001. The bottom row depicts the Euclidean distance maps for the three considered (convex) substrates.

### 2.3. Distance to *κ*_2_ < 0 characterizes spatial cell patterning

We hypothesized that the presence of cells on the convex regions with *κ*_2_ ≥ 0, which was more apparent on day 8 than on day 5, was caused by a collective crowding effect, whereby cells expand from preferentially curved regions into less favourable territory. To investigate this, we created distance maps, quantifying the shortest distance to a region with *κ*_2_ < 0 for every point on the substrate. For the convex cylinder, unduloid, and spherical substrates, which all contain substantial regions with *κ*_2_ ≥ 0, we observed that the distance maps closely resemble the spatial distribution of cells (Figure 2b,e). This observation was quantified by plotting the normalized intensity versus the distance value (termed 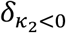), clearly demonstrating a reduction in intensity for increasing distance (Figure 2e). The effect is particularly clear on day 5, where the intensity rapidly drops off to zero in all three cases. On day 8, the rate of intensity reduction is lower, as cells have collectively ventured onto all regions of the substrate, albeit at a lower density for higher 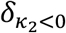. We, therefore, found that, at long enough time scales, the cells can collectively conquer curvatures that are not initially attractive. In this regard, it is not the instantaneous curvature that governs cell patterning, but rather the presence of a region with *κ*_2_ < 0 in the vicinity.

### 2.4. Cells collectively detach from large regions with *κ*_2_ < 0

Closer inspection of the image stacks on the concave substrates revealed that cells do not uniformly fill the concavities, but form suspended cell sheets that span the entire curved region while remaining anchored to the substrate through cell bridges (Figure 3a and Supplementary Movie 2). As demonstrated on a spherical substrate (Figure 3b), the establishment of a detached sheet begins with the individual exploration and spreading of cells in the spherical well. After 5 days, the cell density is high enough for the cells to link up and exert tensile forces to each other, enabling them to lift off the substrate and form bridges. After 8 days, the bridging cells have coalesced into sheets that span the entire concavity, while cell bridges underneath the sheet form anchors to the substrate. These phenomena were not exclusive to the spherical substrates but were observed in all concave substrates after 8 days, across the entire substrate length (Figure 3c). Curvature-induced cell bridging has also been observed in individual mesenchymal stromal cells (MSCs) ^8,33^, and cell sheet detachment has been reported in smooth muscle cells ^34^ and cardiomyocytes ^35^ seeded in microgrooves, though along shorter lengths than we have observed.

**Figure 3:**
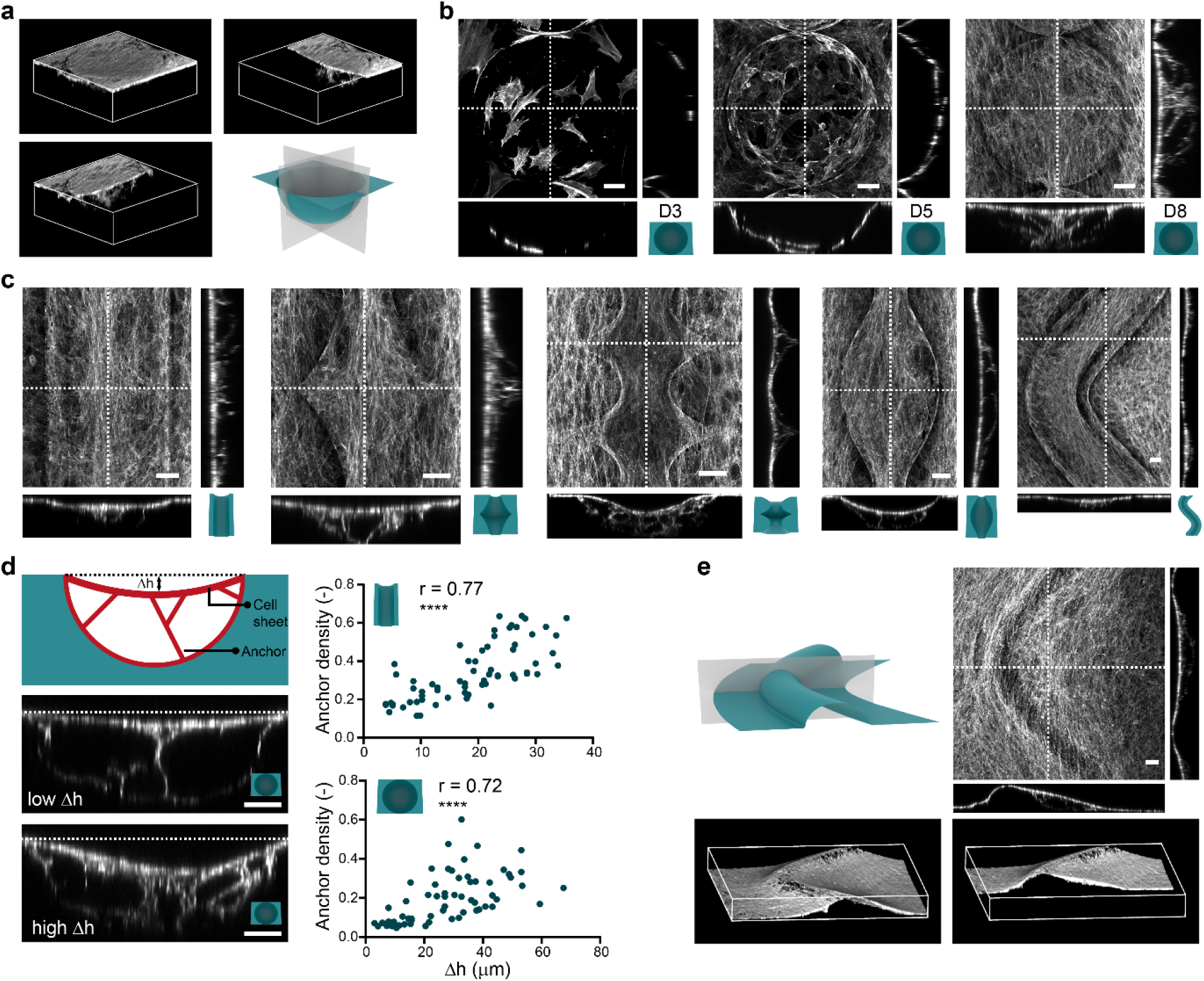
Curvature-induced cell bridging and cell sheet detachment. a) Cell sheet bridging over a concave spherical substrate (*K* > 0, *H* < 0). The 3D reconstruction in the top left panel displays the full coverage of the spherical well, while the cut-away views demonstrate the anchoring cell bridges that connect the sheet to the substrate (see also Supplementary Movie 2). b) Maximum intensity projections and cross-sectional views on spherical wells at days 3, 5, and 8, showing the temporal evolution of bridge formation and sheet detachment. On day 3 (left panel), the cells spread and individually explore the spherical substrate. On day 5 (middle panel), the cells start to link up and form bridges, thereby detaching their body from the substrate. On day 8 (right panel), the cells form detached sheets that completely cover the concave well and are anchored to the substrate through clear cell bridges. c) Maximum intensity projections and cross-sectional views, showing the formation of cell sheets on the other concave substrates. From left to right: cylinder, pseudospheres, catenoids, unduloid, and sinusoidal cylinder. d) Quantification of cell sheet displacement (Δ*h*) in relation to the anchor density for concave cylindrical substrates (top graph) and concave spherical substrates (bottom graph). The *r-value* indicates the Spearman’s correlation coefficient; **** *p* < 0.0001. The data from the cells treated with TGF-β and blebbistatin are also included in these graphs. e) Cell sheet detachment also occurs at local concavities on predominantly convex substrates, as demonstrated for the convex sinusoidal cylinder. The top right panel displays a maximum intensity projection and cross-sectional views, while the bottom panels display 3D reconstructions (full view on the left, cut-away view on the right). All scale bars represent 50 μm.

When evaluating cell sheet detachment across specimens, we observed a variation in the vertical sheet displacement (Δ*h* in Figure 3d). To discern whether this was related to the presence of anchors, we calculated Δ*h* and the anchor density below the sheet for standard experiments on spheres and cylinders, as well as for the experiments with up- or downregulated contractility (see Section 2.6). As expected, we observed a positive correlation between Δ*h* and the anchor density (Spearman’s *r* = 0.77 for the cylinders and Spearman’s *r* = 0.72 for the spheres). We also looked for evidence of cell sheet detachment on the convexly curved substrates. We observed cell sheet detachment at the convex-to-planar transition (*κ*_2_ < 0) on all convex structures, though most notably on the sinusoidal cylinder (Figure 3e and Supplementary Figure 2). Interestingly, we found that sheet detachment is much more pronounced at the concave side of a substrate bend, exhibiting a detached sheet that departs from the top of the substrate and is suspended over a distance that is 4 times longer than at the convex side of the bend (Figure 3e and Supplementary Movie 5). At the concave side, the cells collectively sense the concavity of the transition region (*κ*_2_ < 0) as well as the overall concavity of the substrate (*κ*_1_ < 0). The combination of these curvatures (*K* > 0, *H* < 0) appears to stimulate the formation of a substantial detached cell sheet, suspended over a large region of the planar surroundings.

### 2.5. Curvature induces collective stress fibre orientation

We next asked how curvature affects the collective orientation of stress fibres (SF) on our substrates (Figure 4a). While curvature-induced orientation has been observed in individual cells ^13,36^, cells in monolayers and in developing tissues have shown to cooperatively sense weaker curvature fields ^15,30^. From a differential geometric perspective, it is interesting to compare the orientation of SF to the principal directions of the curved substrates, which are the directions along which *κ*_1_ and *κ*_2_ occur (Figure 4b and Supplementary Figure 3). The cells with pronounced SF have been previously found to align along the direction of minimum principal curvature 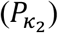, which is often attributed to the tendency to minimize the bending energy of SF ^13,20^. We calculated the degree of alignment (DA, see Materials and Methods) between the SF and 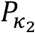 on the convex substrates. The spherical substrates were excluded, since the principal directions are not defined for spheres (*κ*_1_ = *κ*_2_). The strongest DA was observed on the cylinders, showing the collective orientation of SF along the zero-curvature direction (*κ*_2_ = 0) (Figure 4c). While the cells were also found to align well with 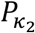 on the unduloid, this was not found to be the case for the pseudospheres and catenoids. On those saddle-shaped substrates, the DA distributions indicate a lower overall alignment with 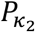, yet the peaks at DA = 0 suggest some alignment with 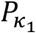 (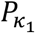 and 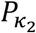 are orthogonal). Interestingly, the cells on the sinusoidal-cylinder were found to collectively align, yet with some deviation from the principal direction 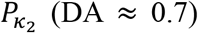. We attribute this deviation to a bypassing effect, whereby SF follow the sinusoidal orientation of the substrate to some extent, but exhibit a collective resistance to change their orientation in response to the alternating curvatures.

**Figure 4:**
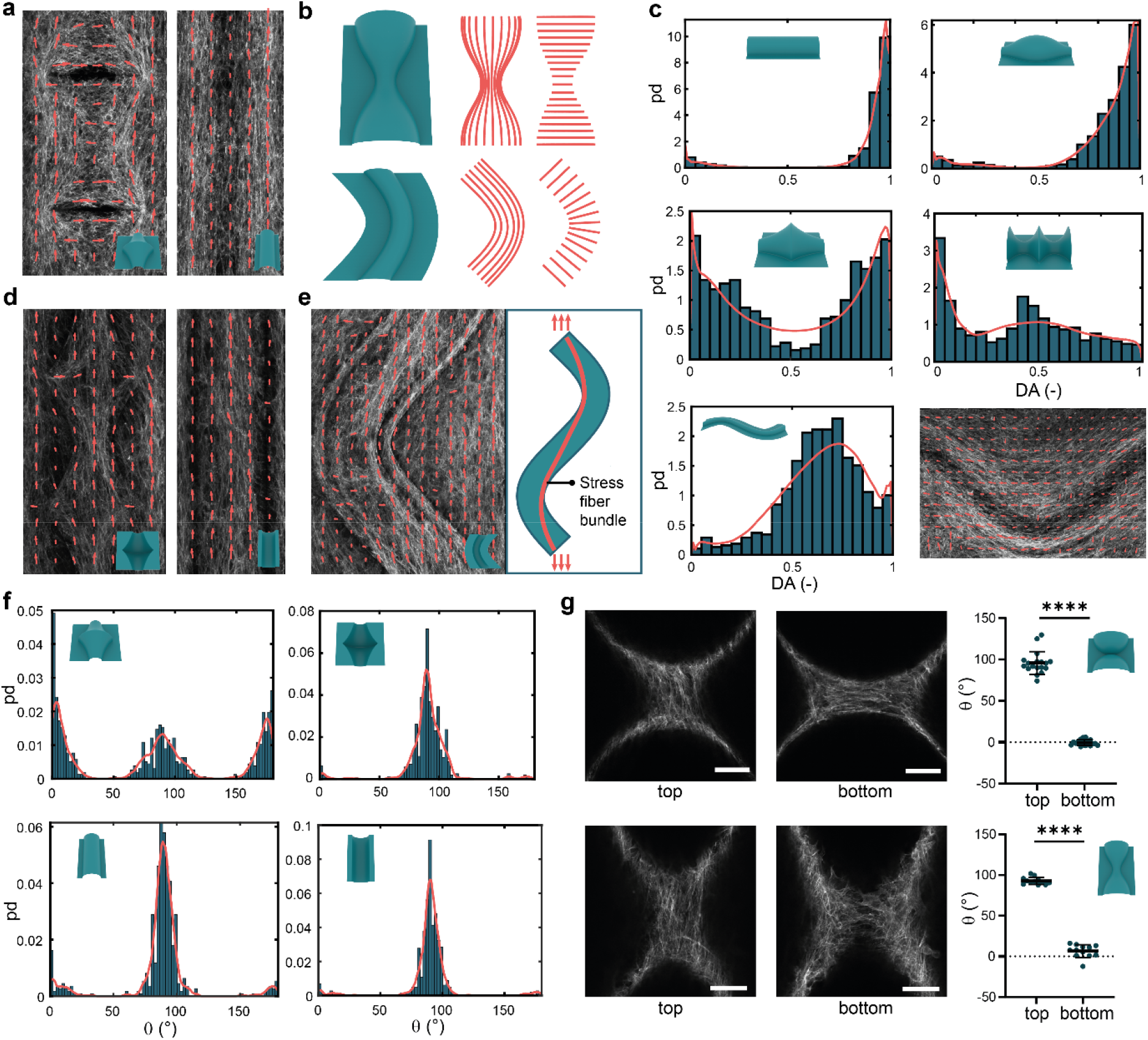
Collective stress fibre orientation on curved substrates. a) Local dominant stress fibre orientations at day 8 on the convex pseudospheres (left) and convex cylinder (right), computed for superpixels containing 80×80 pixels. b) Illustrations of the principal directions on some portions of the convex unduloid (top) and convex sinusoidal cylinder (bottom). The left and right set of lines represent the principal directions corresponding to the minimum (*κ*_2_) and maximum (*κ*_1_) principal curvatures, respectively. c) Probability density distributions (PD) of the degree of alignment (DA) of the SF with respect to the first principal direction (corresponding to *κ*_2_) on the convex substrates. When DA=1, the stress fibres are perfectly aligned with the first principal direction. Data are obtained from all the specimens at day 8 (*n* > 3), using superpixels of 80×80 pixels. The red lines represent the Epanechnikov kernel density estimates. The inset figure on the bottom right displays the local orientation of SF on the convex sinusoidal cylinder at day 8, displaying collective deviation from the first principal direction. d) Orientation of SF at day 8 on the concave pseudospheres (left) and concave cylinder (right), computed for superpixels of 80×80 pixels. e) Orientation of SF at day 8 on the concave sinusoidal cylinder. The right panel schematically illustrates the presence of the central SF bundles that display a lower-amplitude wave, effectively smoothing the geometry of the original sinusoidal substrate. f) PD distributions of the local orientation of SF (with respect to the horizontal axis) on the convex and concave pseudospheres (top left and top right, respectively), and on the convex and concave cylinders (bottom left and bottom right, respectively). The data were obtained from all the specimens at day 8, using superpixels of 80×80 pixels. The red lines represent the Epanechnikov kernel density estimates. g) SF subpopulations on the convex spheres (top row) and convex unduloids (bottom row), displaying orthogonal orientations. The top layer of SF (left) orients along the longitudinal direction (90°), while the bottom layer (right) adopts a horizontal orientation (0°). The graphs on the right display the computed orientations of SF. The data were obtained from three specimens in each case, at multiple locations along the periodic substrates. Paired two-tailed t-tests: **** *p* < 0.0001. Scale bars represent 50 μm.

On the concave substrates, where the cells coalesced in detached sheets, we also observed the collective alignment of SF. In the channel-like substrates (*i.e.*, cylinder, unduloid, catenoids, and pseudospheres), we found a strong longitudinal preference, typically exemplified by the presence of a central SF bundle (Figure 4d,f). This longitudinal preference was attributed to a confinement effect ^37^, where cell crowding in the detached sheet induces collective alignment after 8 days. Indeed, we did not observe longitudinal alignment in the concavities after 5 days, when cells were forming randomly-oriented local bridges (Supplementary Figure 2b). Considering the concave sinusoidal cylinder, we observed SF bundles that trace a lower-amplitude sinusoidal path than the original substrate (Figure 4e). This path is reminiscent of the shape that a tensioned string confined to a sinusoidal channel would adopt, implying a role for actomyosin contractility in the collective orientation of SF.

Finally, we investigated the SF orientation in the regions where the superimposed layers of cells were observed, such as on the saddle-shaped region of the convex unduloid or at the saddle-shaped transition between convex spheres. Interestingly, we found distinct SF subpopulations with orthogonal orientations on both substrates (Figure 4g). In the lower focal planes, horizontally-oriented SF were observed, while the top focal planes displayed vertically-oriented SF. In a different study, orthogonally-oriented SF have been observed within individual cells on saddle shapes ^18^. It was postulated that SF above the nucleus preferentially align in the concave direction to minimize bending, while SF below the nucleus align in the convex direction to support cell migration. Based on these observations, we hypothesize that lower SFs align in the convex direction when cells migrate onto the substrate from both sides and form mechanical connections. Once these regions have been conquered, new cells can align in the concave direction to minimize bending.

### 2.6. Contractility and differentiation perturbation affect curvature guidance

We explored the role of contractility on the collective organization in our complex, curved landscapes. We enhanced cell contractility with transforming growth factor-β (TGF-β) and downregulated contractility using myosin-II-inhibiting blebbistatin. In general, the cells with perturbed contractility patterned the curved substrates similarly to the unperturbed cases, yet exhibited more- or less-pronounced SF in response to TGF-β and blebbistatin, respectively (Figure 5a and Supplementary Figure 4). However, we found that the cells treated with blebbistatin did not cover the unfavourably curved regions (*κ*_2_ > 0) as well as unperturbed cells (Figure 5b).

**Figure 5:**
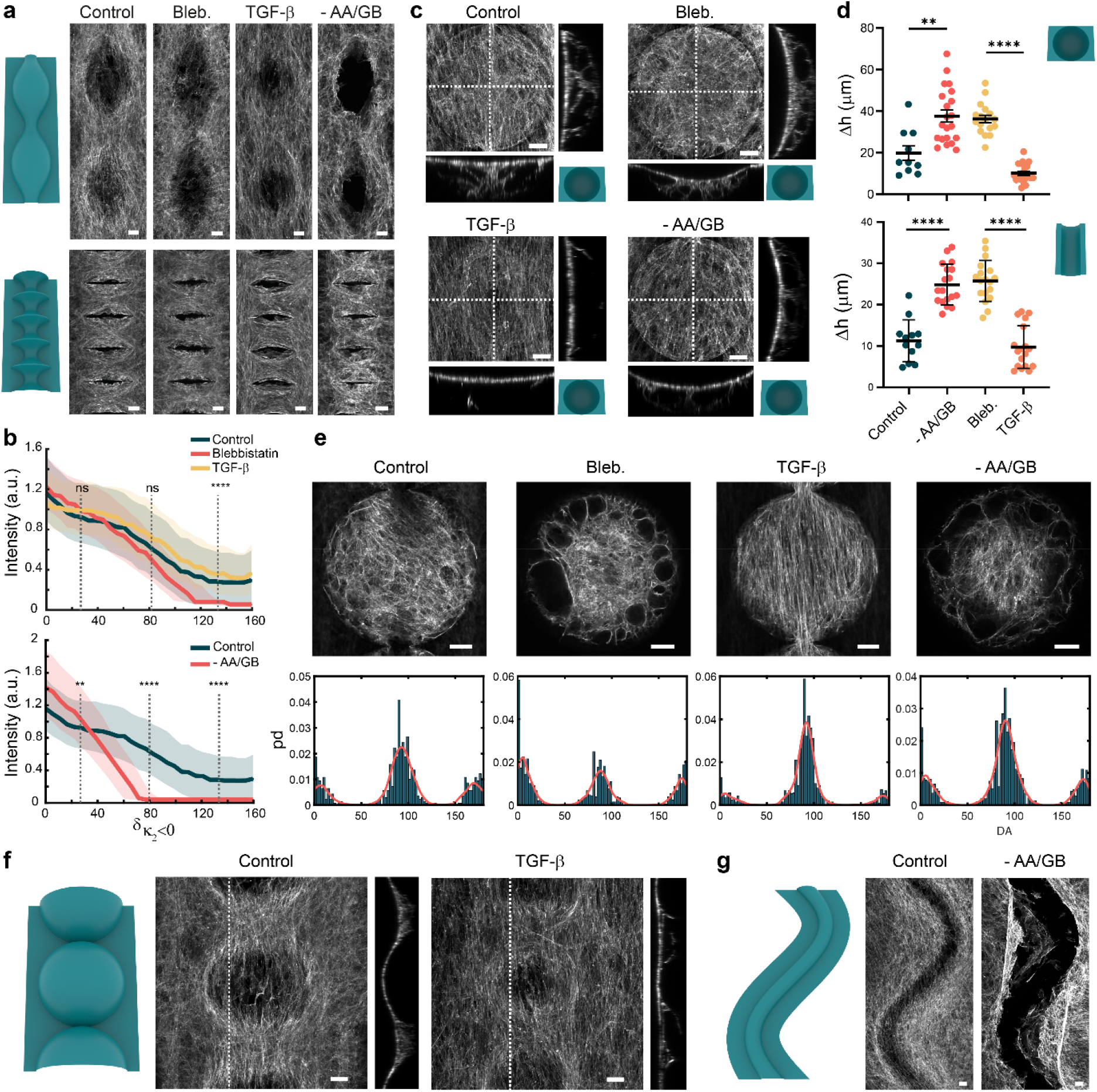
Effect of contractility perturbation and differentiation inhibition on the curvature-induced organization. a) Representative maximum intensity projections (actin) on the convex unduloids (top) and catenoids (bottom), displaying the effects of contractility inhibition (Bleb.), contractility enhancement (TGF-β), and differentiation inhibition (-AA/GB). b) Intensity reduction as a function of 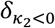. The top chart demonstrates the effects of contractility perturbation. The Kruskal-Wallis test: **** *p* < 0.0001. The bottom chart demonstrates the effects of differentiation inhibition. The Mann-Whitney test: ** *p* < 0.01, **** *p* < 0.0001. The solid lines represent the median values while the shaded areas represent the interquartile range. c) Effect of contractility perturbation and differentiation inhibition on cell sheet detachment over concave hemispheres at day 8. d) Quantification of cell sheet displacement (Δ*h*) due to contractility perturbation or differentiation inhibition. e) Effects of contractility perturbation and differentiation inhibition on the morphology and orientation of SF. The top row displays representative slices through the confocal z-stacks, while the bottom row displays the PD of the SF for all the specimens in each category. f) Contractility enhancement affects bridge formation on the convex spherical substrates. A stronger cell sheet is observed in the specimens treated with TGF-β. g) Differentiation inhibition causes cell sheet detachment on the convex sinusoidal cylinders in some specimens. The cells collectively pull away from the concave bend of the substrate. All scale bars represent 50 μm.

Cell contractility is the driving force behind the formation of individual or multicellular bridges. Indeed, we found that perturbing contractility affects the morphology of the detached cell sheets and anchoring bridges on the concave substrates (Figure 5c and Supplementary Movies 3-4). Enhancing contractility results in lower sheet displacement and fewer bridges as compared to the cases where contractility is inhibited (Figure 5d), implying that higher cell contractility translates to a higher overall tension in the suspended sheets. When considering specific focal planes within the cell sheets of the different cases, we also observed different SF morphologies and orientations. As compared to the unperturbed case, the cells treated with blebbistatin are more dendritic-like and form many small anchoring bridges that adopt circular configurations, while TGF-β induces strong SF with a more pronounced longitudinal alignment (Figure 5e). The effects of collectively enhanced contractility were also apparent on the convex substrates, where strong cell sheets were observed that bridge the underlying curved structures. On the spherical substrates, for example, the cells cultured under normal conditions form modest bridges in between the spheres, while contractility-enhanced cells form much stronger cell sheets that remain almost planar and seem to avoid the underlying curvature (Figure 5f). These results are in line with previous studies at the cell and tissue scales ^19,30^, and underpin the important role of individual cell contractility as a driving force for the collective organization in varying curvature fields.

We also stained cells for runt-related transcription factor 2 (RUNX2), a marker for osteoblast differentiation. We observed that RUNX2 was clearly expressed in the cells in the control group, but also in specimens with up- or downregulated contractility. When considering RUNX2 expression on the spherical and saddle-shaped regions of the convex unduloid (constant mean curvature), we consistently observed significantly higher RUNX2 intensities on the saddle-shaped (*κ*_2_ < 0) regions as opposed to the spherical (*κ*_2_ > 0) regions (Supplementary Figure 6). This points towards a role for curvature in the regulation of cell differentiation, which has been reported by others in MSCs ^19,33^.

The osteogenic culture medium that was used during our experiments contained ascorbic acid (AA) and β-glycerophosphate (GB), which support the development of mineralized extracellular matrix (ECM) and promote osteoblast differentiation ^38^. AA induces the secretion of type I collagen (Col1) in the ECM, while GB works synergistically with AA and acts as a phosphate source for mineralization ^38–40^. To study how collective curvature-guidance is affected by the inhibition of ECM development, we performed experiments with a culture medium deprived of AA and GB ^41^. While the cells cultured in this medium generally exhibited similar curvature-induced organization, we observed significantly lower degrees of cell coverage on unfavourably-curved regions (*κ*_2_ > 0) (Figure 5a,b). On the concave structures, we found that AA/GB deprivation resulted in weaker cell sheets, displaying similar SF morphologies and Δ*h* as compared to the cells treated with blebbistatin (Figure 5c-e). Interestingly, we observed that the cells cultured in non-osteogenic medium in some cases collectively pulled away from the concave bends of the convex sinusoidal cylinders, a phenomenon that was never observed in the cells cultured in osteogenic medium (Figure 5f). These results indicate that the development of ECM, induced by osteogenic medium, provides a reinforcing scaffold that enables the cells to collectively conquer unfavourable curvatures within our complex landscapes.

## 3. Discussion

We have demonstrated the collective organization of preosteoblasts in cell-scale, varying-curvature landscapes. By designing mathematically-defined surface families with controlled curvature variations and by leveraging high-resolution free-form fabrication, we micro-engineered substrates that cover a wide range of Euclidean, hyperbolic, and elliptical geometries. This broad curvature spectrum enabled us to systematically study collective curvature-guidance and unify recent findings within a geometry-centred framework. We found that cells preferentially pattern regions with at least one concave direction (*i.e.*, regions where the minimum principal curvature is negative, *κ*_2_ < 0), while curved regions with *κ*_2_ ≥ 0 are generally avoided. Previous studies have also reported preferences for local concavities, which has been attributed to a more relaxed stress configuration in the concavities ^19,29,33^. However, those observations were typically made from a 2D perspective or without addressing the formal mathematical framework of surface curvature. Specifically, curvature guidance has often been studied by considering cell behaviour on convex or concave substrates, typically spheres or cylinders, without addressing the specific mean or Gaussian curvatures. Moreover, hyperbolic substrates, containing both concave and convex directions, have received little attention until recently ^14,30^, despite their high physiological relevance ^8^. Our results show that one concave direction in a specific region is sufficient for cells to preferentially cover that region. This includes not only all hyperbolic geometries (*K* < 0) but also elliptical (*K* > 0) and Euclidean (*K* = 0) regions where the mean curvature is negative, such as the convex-to-planar transitions that have been typically ignored in previous studies. In this regard, there is no single mean or Gaussian curvature that could be pinpointed as ideal for cell patterning, but rather a spectrum of shapes where *κ*_2_ < 0. Despite a general preference for *κ*_2_ < 0, we found that cells can eventually conquer unfavourably-curved regions through cooperative action, provided that the distance to favourably-curved regions is not too large. However, this ability to venture onto curved regions with *κ*_2_ ≥ 0 reduces when cell contractility or ECM production is impaired.

Our widespread observations of cell bridging and cell sheet detachment, including observations with blebbistatin and TGF-β, further illustrate the crucial role of cell contractility and the ability of the cells to collectively override the geometrical cues imposed by the substrate. However, an important parameter governing these phenomena is the cell adhesion to the substrate. Our substrates were functionalized by fibronectin (FN) adsorption, facilitating integrin-mediated adhesion. However, different results might be obtained when the FN is covalently bonded to the substrate ^30^. In such cases, cell sheet detachment may be reduced or avoided ^34^.

Our results on the curvature-induced collective orientation of SF are in line with the biophysical arguments that SF-dominated cells align in directions that minimize SF bending ^13,20^. Indeed, we find that cells on convex cylinders or unduloids align well with the direction of minimum principal curvature. On the convex hyperbolic substrates (where *H* > 0), however, we found that cells show less uniform alignment, and that a substantial portion of SF align along a locally convex direction. Moreover, we observed orthogonally-oriented SF subpopulations on some local saddles and confinement-induced longitudinal orientation of SF in detached cell sheets. In general, we concluded that substrate curvature, indeed, affects SF orientation, but that cell interactions, mediated by contractility and ECM production, result in a collective resistance to local variations in the underlying curvature.

Taken together, our results underpin the importance of local, cell-scale geometrical cues on the emergent organization of bone-like cells. In particular, these findings emphasize the role of multicellular cooperation, enabling cells to collectively conquer unfavourably-curved regions or alter their local environment through collective detachment. However, cells are typically exposed to several other biophysical cues *in vivo*, such as stiffness gradients ^42^ or nanotopographies ^6^. In this regard, it would be interesting to explore the collective cell organization in a tailored multi-cue environment to unravel the dominant cues and potential crosstalk ^14^. Our results could ultimately inspire the design of tissue engineering scaffolds ^10^. Based on our findings, one could argue that scaffolds with substantial regions with *κ*_2_ < 0 are preferred. In scaffolds based on cylindrical strut networks, which have often been proposed ^43–46^, such regions occur at the intersections of struts. Indeed, tissue formation has been found to initiate from those locations *in vivo* ^47^. Alternatively, one could consider hyperbolic sheet-based scaffolds, such as those based on triply periodic minimal surfaces (TPMS), which have *κ*_2_ ≤ 0 at every point. However, developing geometrically-optimized scaffold designs requires further investigation into the intricacies of cell-geometry interaction, likely involving computational studies that take geometry explicitly into consideration ^48^. Nevertheless, fuelled by rapid advances in high-resolution free-form fabrication, we anticipate exciting avenues for geometric control of cells and tissues, relying on surface curvature as the language of shape.

## Acknowledgements

The first author is thankful to Milica Dostanic and Paul Motreuil-Ragot for initial discussions and assessment of the PDMS moulding. We are also thankful to Prof. Dr. Urs Staufer from the Precision and Microsystems Engineering department at TU Delft, for enabling the use of the Nanoscribe GT2 device. The research leading to these results has received funding from the European Research Council under ERC grant agreement no. [677575].

## Data availability

The codes and data used to obtain these results can be made available upon reasonable request from the first author.

## 4. Materials & Methods

### Design of curved substrates

The curved substrates were designed in MATLAB (MATLAB 2018b, Mathworks, Natick, MA, USA) and SolidWorks (Dassault Systèmes, Vélizy-Villacoublay, France). For the axisymmetric substrates (*i.e.*, unduloid, catenoid, sphere, pseudosphere, and cylinder), the generatrix curves were generated in MATLAB using the parametrizations provided in Supplementary Note 1. The curves were then imported into SolidWorks and π-revolutions around the central axis were generated, resulting in convex half-surfaces of revolution. Next, end-caps and a rectangular bottom layer (with 20 μm thickness) were added to convert the surfaces to printable solids. The sinusoidal cylinder substrate was directly designed in SolidWorks, by sweeping a hemi-circular cross-section along a sinusoidal guiding curve. The corresponding substrate designs were exported in the STL format, to prepare them for the printing process.

### Fabrication and functionalization of PDMS substrates

Mould masters were fabricated using a two-photon lithography 3D printer (Nanoscribe GT2, Nanoscribe GmbH, Karlsruhe, Germany). A silicon substrate was cleaned with isopropanol (IPA, Merck KGaA, Darmstadt, Germany) and was treated with oxygen plasma (Femto, Diener electronic GmbH + Co. KG, Ebhausen, Germany) to improve its adhesion. A droplet of IP-S acrylate-based resin (Nanoscribe GmbH) was drop cast onto the substrate. Four sets of six convex master substrates were 3D printed on the silicon substrate using a 25X objective (NA = 0.8), 0.5 μm hatching, 0.5 μm slicing, 50 mW nominal laser power, and 50 mm/s scanning speed. To save time, an internal support scaffold was written instead of a solid block. The structures were developed in propylene glycol methyl ether acetate (PGMEA, Merck KGaA, Darmstadt, Germany) for 25 minutes followed by 5 minutes of treatment with isopropyl alcohol (IPA) and blow drying using a filtered air gun.

To perform the moulding, the master was first placed in a vacuum desiccator beside a glass petri dish containing a droplet of trichloro(1H,1H,2H,2H-perfluorooctyl)-silane (Merck KGaA, Darmstadt, Germany), which subsequently coated the surface of the master with a hydrophobic layer allowing easy future peeling of the polydimethylsiloxane (PDMS) copy. PDMS (Sylgard 184, Dow Inc., Midland, MI, U.S.A.) mixed with the curing agent at a weight ratio of 10:1 was mixed thoroughly, drop-cast on the master, desiccated in vacuum for 30 minutes to remove any air bubbles, and cured in an oven at a temperature of 40 °C for 16 hours. The resulting copy was cut out by scalpel and gently peeled off the master. Typically, 7-10 PDMS copies were made with a single master without loss of fidelity. This single-step moulding process resulted in specimens with concave curved substrates (imprints in the PDMS). To obtain the convex counterparts (protrusions), a double moulding step was performed. To this end, a PDMS copy originating from the single-moulding operation was thermally aged in an oven at 100 °C for 48 hours ^49^. This PDMS substrate was consequently used as a new mould for the second moulding stage, which was performed in the same way as before. After curing, the second PDMS could easily be peeled off the PDMS mould, without loss of fidelity. The quality of the single-step and two-step PDMS moulding processes was verified using a scanning electron microscope (SEM, JSM-IT100LA, JEOL, Tokyo, Japan) with a beam energy of 10 kV and a working distance of 12 mm. Prior to SEM imaging, the PDMS substrates were gold sputtered (layer thickness of 5±2 nm) to enhance conductivity. Additionally, the quality of the replica moulding procedures was confirmed through laser confocal scanning (Keyence VK-X 3D scanner, Keyence, Osaka, Japan) using a 20x magnification lens.

In preparation for cell seeding, circular specimens of 8 mm diameter were punched from the PDMS copies. Next, the specimens were sterilized inside an oven at 110 °C for 1 hour. To reduce the inherent hydrophobicity of PDMS and facilitate substrate wetting, the specimens were subsequently treated with oxygen plasma (Femto, Diener electronic GmbH + Co. KG, Ebhausen, Germany) for 3 minutes. Then, the PDMS specimens were transferred to a 48-well plate, washed twice with 10 × PBS, submerged in a solution of 50 μg/ml bovine fibronectin (Sigma-Aldrich, St. Louis, MO, USA), and incubated at 37 °C and 5% CO_2_ for 30 minutes to functionalize the PDMS surface and promote cell adhesion. After the removal of the fibronectin solution, the specimens were thoroughly washed with 10 × PBS.

### Cell seeding and culture

Prior to cell seeding, murine preosteoblasts (MC3T3-E1, Sigma-Aldrich, St. Louis, MO, USA) were cultured for 7 days in minimum essential medium (α-MEM, Sigma-Aldrich) with the addition of 10% fetal bovine serum and 1% penicillin-streptomycin (both from Thermo Fischer Scientific, Waltham, MA, USA). The medium was refreshed every 2 to 3 days. In all the experiments, approximately 5 × 10^3^ cells were seeded on the fibronectin-coated PDMS specimens in 250 μl culture medium, which were then cultured for up to 8 days at 37 °C and 5% CO_2_ with the medium being refreshed every 2 to 3 days. To induce osteogenic differentiation, the culture medium was supplemented with 4 mM β-glycerophosphate and 50 μg/ml ascorbic acid (both from Sigma-Aldrich), starting from day 3. In the experiments where differentiation was inhibited, the cells were cultured in the standard culture medium throughout the entire duration of the experiments (*i.e.*, without the addition of ascorbic acid or β-glycerophosphate). In the experiments with enhanced cell contractility, 1 ng/ml of TGF-β3 (Sigma-Aldrich) was added to the culture medium on day 3 and this concentration was maintained throughout the remainder of the experiments. To inhibit cell contractility, the culture medium was supplemented with 10 μM of blebbistatin (Sigma-Aldrich), which was also maintained throughout the remainder of the experiments.

### Immunostaining

Immunostaining was performed at different time points (days 3, 5, and 8). The specimens were washed twice in 10 × PBS and fixated in 4% formaldehyde/PBS for 15 minutes at room temperature. Next, the specimens were washed with 1 × PBS and the cells were permeabilized in 0.5% Triton/PBS at 4 °C for 5 minutes, followed by incubation in 1% BSA/PBS at 37 °C for 5 minutes. To stain for F-actin, the specimens were incubated in 1% BSA/PBS with rhodamine phalloidin (1:1000, Thermo Fischer Scientific). Afterwards, the specimens were washed 3 times for 5 minutes with 0.5% Tween/PBS at room temperature, followed by washing for 5 minutes with 1 × PBS at room temperature. The specimens were subsequently mounted in a glass-bottom dish using a droplet of ProLong Gold antifade reagent with 4’,6-diamidino-2-phenylindole (DAPI, Thermo Fischer Scientific) to stain the chromatin cargo in the nuclei. The specimens that were also stained for RUNX2 followed a similar protocol, involving a first incubation step with anti-RUNX2 primary antibody (Abcam, Cambridge, UK) followed by Tween/PBS washing and a second incubation step with Alexa Fluor 488 conjugated secondary antibody (Thermo Fischer Scientific).

### Confocal imaging

Fluorescence confocal laser scanning microscopy (CLSM) was performed using a Nikon Eclipse Ti inverted confocal microscope (Nikon, Tokyo, Japan) with a Nikon Plan Apochromat λ 10x objective (0.45 NA). The images were acquired using 2 or 3 laser lines with excitation wavelengths of 405 nm (DAPI), 488 nm (Alexa Fluor 488), and 561 nm (Rhodamine-phalloidin) with the detection windows set accordingly. Z-stacks were obtained at an *xy*-resolution of 0.60 × 0.60 μm and a z-spacing of 1 μm (for specific cases) to 5 μm (nominal cases). The acquisition in the different channels was performed sequentially to minimize inter-channel cross-talk.

### Preparation of stack projections and curvature maps

Maximum intensity projections were obtained from the image stacks using Fiji ^50^. A custom image registration script in MATLAB was used to select a rectangular region of interest (ROI), defined by the bottom layer of the curved substrates, and crop the image to the ROI. The resulting image was subsequently rotated to align the rectangular ROI in the vertical direction. To assess the relationship between the confocal image data and the curvature of the underlying substrate, curvature maps were created in MATLAB using the parametrizations provided in Supplementary Note 1. The transition region between the curved substrates and the flat surroundings was assigned a radius of curvature of 15 μm to account for the local concavity that this narrow region presents. The curvature maps were defined as pixelated images, matching the resolution of the corresponding confocal images, enabling a pixel-by-pixel comparison of the confocal data and the curvature. The principal curvatures were non-dimensionalised by multiplying with the radius of the spherical substrate (leading to *κ*_1_ = *κ*_2_ = 1 for the convex spherical substrates).

### Actin frequency maps

To create the frequency maps of the actin images (Figure 2b), several images belonging to the same experimental group were split into periodic units and superimposed in MATLAB. First, the intensity was normalized with respect to the mean intensity of the image. All the images within the same group were then summed. Finally, the images were split into periodic units (Figure 2b) and summed again to create the final frequency map.

### Intensity quantification and distance maps

The actin maximum intensity projections were normalized with respect to the mean intensity and were converted to grayscale. Using custom MATLAB scripts, every image was rasterized (downsampled) into a set of “superpixels”, where the intensity of each superpixel is the average of all the pixels it is composed of. Through this rasterized approach, the intensity variations are considered for local neighbourhoods (better corresponding to the scale of the cells), rather than at the individual pixel level. The size of the superpixels is mentioned in figure captions. The distance maps (Figure 2e) were created by binarizing the curvature maps of the minimum principal curvature (*κ*_2_). These binarized curvature maps were used to compute the Euclidean distance transform for which every pixel was assigned a value depending on its distance from the nearest point with a negative minimum principal curvature. To assign a higher weight to the regions with *κ*_2_ > 0 than regions with *κ*_2_ = 0 (corresponding to the observed relative preference of the cells for the latter as opposed to the former regions), the final distance map was defined as the average of two distance maps: a map representing the distance to points with *κ*_2_ ≤ 0 and a map with the distance to points with *κ*_2_ < 0.

### Quantification of cell sheet detachment on concave spheres and cylinders

To quantify the amount of cell sheet displacement and the anchor density (Figures 3d and 5c), the actin z-stacks were first cropped to a square ROI and then resliced from the top with 5 μm spacing in Fiji. Using custom scripts in MATLAB, the middle image of the resliced stack was selected, the detached cell sheet was traced, and the maximum displacement of the sheet with respect to the endpoints of the sheet was quantified (Figure 3d). The anchor density was quantified by masking and binarizing the region below the detached sheet in the resliced stacks. Consequently, the density of the actin voxels in the binarized image stack was determined. The 3D reconstructions of the confocal stacks in Figure 4 and the Supplementary Movies were generated using the open-source Fiji plugin 3Dscript ^51^.

### Quantification of stress fibre orientation

To determine the dominant stress fibre orientation in an ROI, a custom script based on the Fast Fourier Transform (FFT) was implemented, similar to a previously reported method ^41^. Details of the implementation are provided in Supplementary Note 2 and Supplementary Figure 5. Briefly, the approach involved computing the power spectrum of the FFT applied to a grayscale ROI and detecting the orientation of the dominant band of the elevated power values in the spectrum. This orientation was then converted to the dominant orientation in the grayscale image. The orientation analysis was applied to every sub-image in the rasterized images (Figure 4). For all the structures, the maps of both principal directions were created (similar to the curvature maps described before). The DA between the SF and the first principal direction (pd, corresponding to *κ*_2_) was defined as:

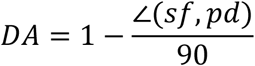

where ∠(*sf, pd*) ∈ (0,90) indicates the angular difference between the SF and the principal direction.

### Statistical analysis

All statistical analyses were performed using GraphPad Prism 8 (GraphPad Software, CA, USA). For all the relevant figures, the type of the data presented, the choice of the statistical tests, and the significance levels are all indicated in the corresponding figure captions.

